# Revision of Indian Bipaliid species with description of a new species, *Bipalium bengalensis* from West Bengal, India (Platyhelminthes: Tricladida: Terricola)

**DOI:** 10.1101/2020.11.08.373076

**Authors:** Somnath Bhakat

## Abstract

A new species of Bipaliid land planarian, *Bipalium bengalensis* is described from Suri, West Bengal, India. The species is jet black in colour without any band or line but with a thin indistinct mid-dorsal groove. Semilunar head margin is pinkish in live condition with numerous eyes on its margin. Body length (BL) ranged from 19.00 to 45.00mm and width varied from 9.59 to 13.16% BL. Position of mouth and gonopore from anterior end ranged from 51.47 to 60.00% BL and 67.40 to 75.00 % BL respectively. Comparisons were made with its Indian as well as Bengal congeners.

Salient features, distribution and biometric data of all the 29 species of Indian Bipaliid land planarians are revised thoroughly. Genus controversy in Bipaliid taxonomy is critically discussed and a proposal of only two genera *Bipalium* and *Humbertium* is suggested.

## Introduction

Terrestrial planarians are relatively species-poor group with only 822 described nomial species worldwide (Winsor 1997, Jones 1998, Sluys 1999, Carbayo et al. 2002). Over 160 species of the genus *Bipalium* Stimpson, 1857 are known in this Asiatic triclad group. This group was also poorly studied due to various reasons namely restricted distribution, nocturnal habit, morphological similarity with earthworm etc. In India, after independence, except a few early workers (Johri 1952, Sharma and Sharma 1977, Rout and Ghose 1979, Ramkrishna and Chauhan 1960), systematic study on this group is lacking. Recently Kawakatsu and Jayashankar (2013) reported a checklist of 27 known species with description of three unidentified *Diversibipalium* spp. from India. In this period of lockdown, I have surveyed my area for different species of animals from invertebrates to vertebrates. On 5th September, 2020, I observed a land planarian on the road side in my locality. Later I studied the animal in detail and in the following days a number of specimens were collected from the same spot. The present paper is based on the description of this new species, *Bipalium bengalensis* n. sp.

In Bipaliid taxonomy (Family: Bipaliidae von Graff, 1896), controversy persists for a long time with new proposal of genus from time to time. Elliot (1948) introduced the genus *Planaria* to describe a few land planarians. Wright (1860) described some hammerhead worm from India and China under the genus *Dunlopea*. Later, all the species of land planarian were put in the genus *Bipalium*. Graff (1899) at first reported three genera of land planarian on the shape of head namely *Bipalium*, *Perocephalus* and *Placocephalus*. In 1998, Kawakatsu et al. erected another genus *Novibipalium* (new “Bipalium”) for those species with reduced or absence of genital papilla. But it draws controversy as in both genera, a set of fold formed the functional penis. Ogren and Sluys (2001) proposed another genus *Humbertium* on the basis of entry of ovovitelloducts to the female atrium. In *Humbertium*, the ducts enter the female atrium anteriorly while posteriorly as in *Bipalium*. Later Kawakatsu et al (2002) created one more genus *Diversibipalium* (the “diverse *Bipalium*”) to include all species whose anatomy of the sexual organ was unknown. In 2013, Kawakatsu and Jayashankar reported four genus viz. *Bipalium*, *Humbertium*, *Novibipalium* and *Diversibipalium* to list 27 Indian Bipaliid species with their distribution in India. The report is incomplete as there is no morphological description or biometric data of the concerned species. So in the present paper, I have revised the Indian Bipaliid species with salient features and biometric data along with their distribution to fill up the lacunae.

## Materials and methods

Live planarians are collected from the moist soil surface of the road side at dawn (5.00 to 5.30 a. m.) from September 5 to September 25, 2020 in the day after rainfall. As the land planarians are nocturnal in habit, after sun rise they hide themselves under stones or soil. The collection site is full of debris, grasses, weeds and other organic matters, an ideal inhabitant of their prey like earthworm, small insects and other soil organisms. The specimens were collected at Suri (87°32’00”, 23° 55’00”N), Birbhum district, West Bengal, India (Fig. 1).

**Fig. 1.**
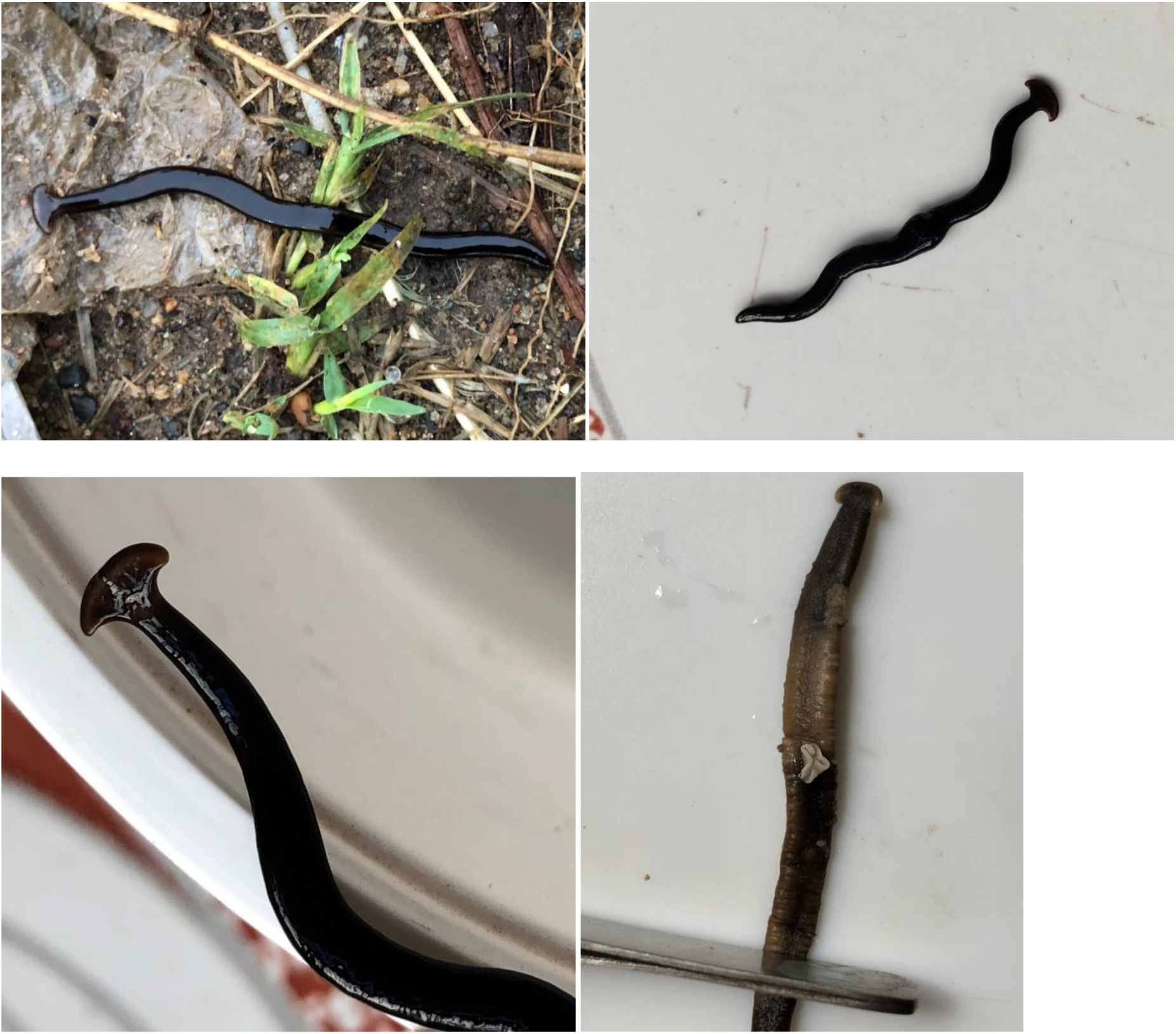
*Bipalium bengalensis* sp. n.in nature, the planarian showing pharyngeal swelling (in the middle of the body), elevated anterior half of the body, protruded pharynx from the dorsal surface (white frill). (clockwise from upper left).

Live animal is transferred in a large watch glass containing 25 – 30% NaCl solution. Within a few minutes, the animal died and then it is transferred to distilled water for washing. Finally the specimen is transferred to 70% alcohol for preservation. In salt solution, a very thin white mucous layer is separated from the body. In this technique, the colour and shape of the body remain intact.

Measurements of each specimen were taken within two days of preservation. All the measurements were made with a digital dial caliper to the nearest 0.1mm. Different measurements of body parts are presented as percentage of body length (BL) while width of sole and head rim is given as percentage of body width (BW) and head breadth (HB) respectively. Mean and standard deviation (SD) of each parameter was calculated separately. Formaldehyde (4%) preserved old specimens (at least two years old) were also measured along with the fresh specimens.

For revision work, information was collected from literature and Tricladida Database.

## Results

### Description of new species

#### Taxonomy

Phylum: Platyhelminthes Claus, 1887
Class: Turbellaria Ehrenberg, 1831
Order: Tricladida Lang, 1884
Family: Bipaliidae von Graff, 1896
Type species: *Bipalium bengalensis* sp. n.

#### Type material

##### Holotype

05. IX. 2020.; Suri (Vivekanandapally) (87.5151oE, 23.9146oN), Birbhum district, West Bengal, India; 28.4mm BL; Zoological Museum, Department of Zoology, Rampurhat College, Rampurhat-731224, Birbhum district, West Bengal, India; Collecter: Somnath Bhakat.

##### Paratype

Same locality; 19.00-45.00mm BL (n= 11); September 5-25, 2020; all other details are same as holotype.

### Description: (Fig.1) (Table 1)

Elongated bilaterally symmetrical and dorsoventrally flattened body is with a semilunar head. The anterior portion of the body connecting the head is narrow to form neck, gradually broadens to reach maximum width at mouth region. In some specimen, mouth region is more expanded than normal, form pharyngeal swelling (Fig. 1).The posterior most part of the body is narrow with blunt tip. Head plate is semilunar in shape with numerous small eyes limited to the margin of the head, form a rim (0.7 – 1mm width). In live condition, when the planaria search for a suitable surface, a wavy movement is observed in the peripheral margin of the head. In normal condition, the head is slightly elevated from the surface to investigate any object but in critical condition, the worm can elevated/raised half of its body length to attach with a suitable surface (Fig. 1). Mouth opens beyond 50% and the gonopore in the 3/4th of the body from anterior end. In some specimen, another yellowish circular spot is present followed by gonopore. Elevated creeping sole begins at the base of the head, extending to the tip of the posterior end where it is wider to form a dumbbell (Fig. 2).

**Table 1.**
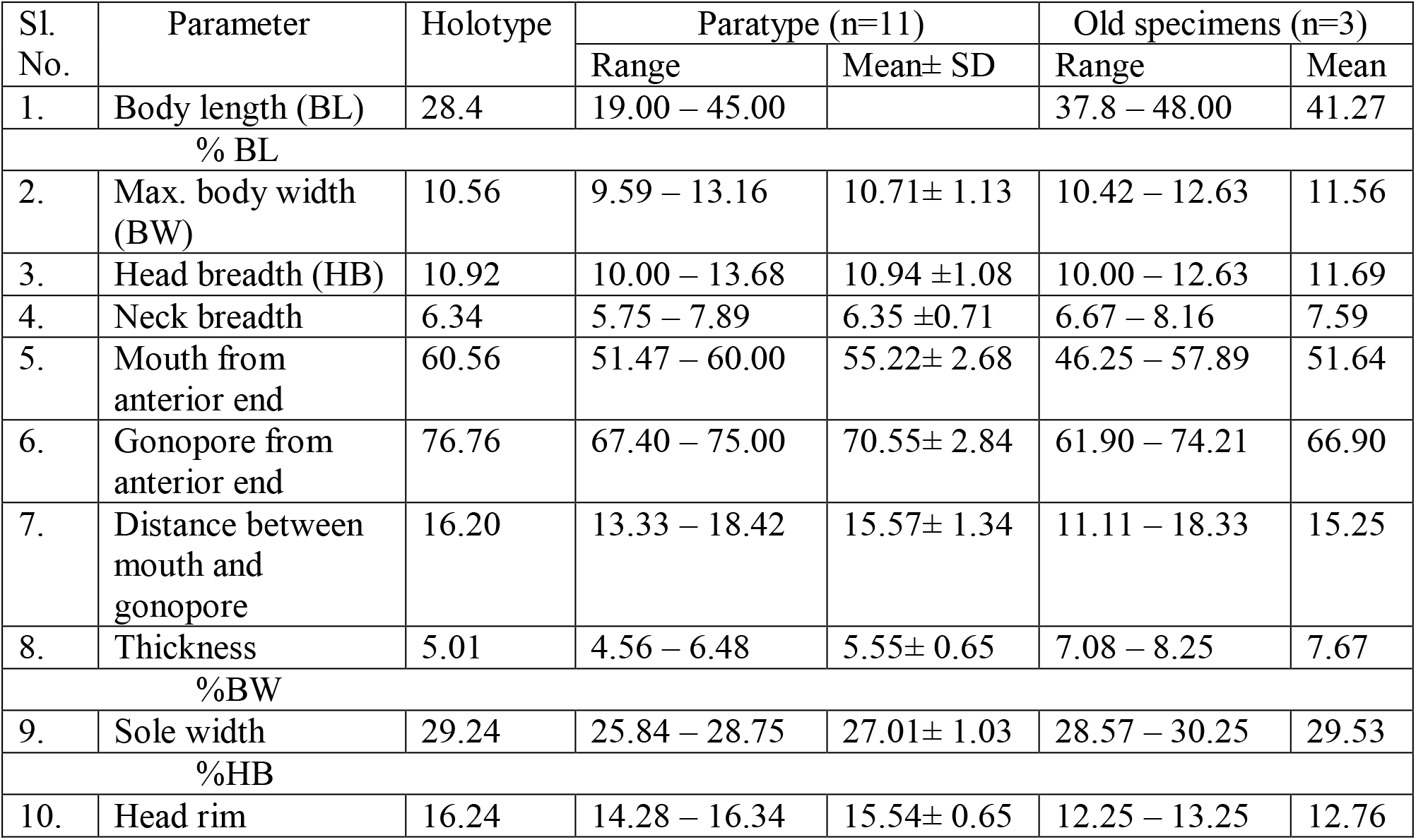
Biometric data of *Bipalium bengalensis* n. sp. (BL in mm.).

**Fig. 2.**
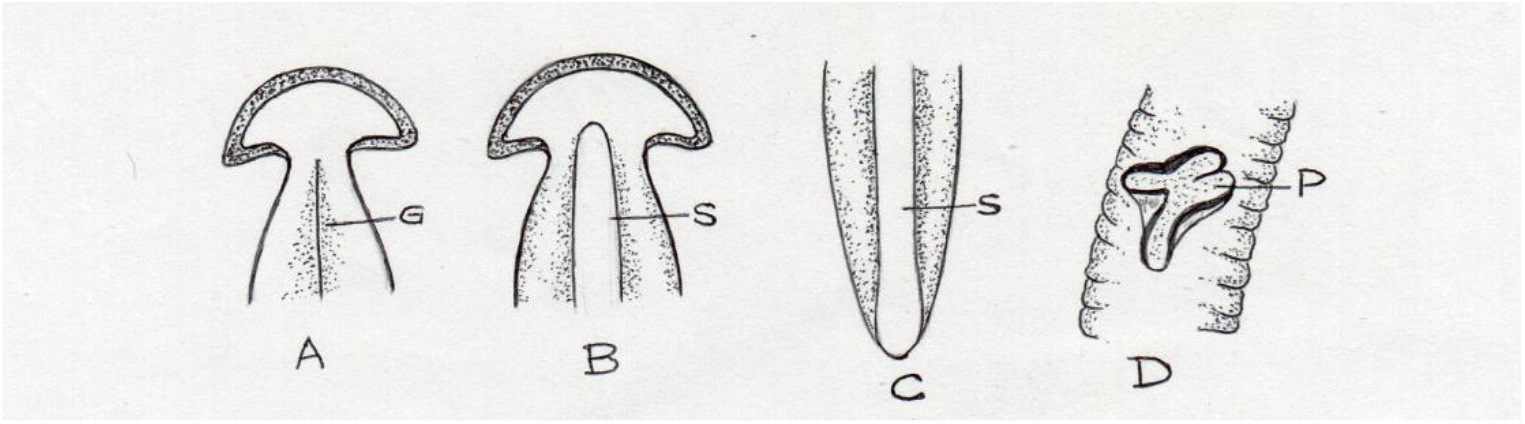
*Bipalium begalensis* sp. n. A. Anterior portion showing head and median groove (G), B. Ventral portion showing sole (S), C. Posterior end showing dumbbell sole (S), D. Enlarged protruded pharynx (P) with wrinkled body margin (in preserved specimen).

Colour of the dorsal surface is jet black with a narrow longitudinal groove in the middle. In live condition, the head plate is pale black with pinkish rim. Auricle of semilunar head plate is pale in colour. The ventral surface is pale dark with yellowish white creeping sole in the middle (Fig. 2).

In the formaldehyde preserved old specimen (more than two years old), the body became more thick, wrinkled and tough. The dorsal surface is decolourised and turned into yellowish brown with tinge of black though anterior 1/4^th^ is black mottling. Head is blackish with yellowish margin. Ventral surface of head is whitish brown. Pale yellowish ventral is with a distinct elevated whitish sole in the middle. In one specimen, pharynx with white thick margin, is protruded through the mouth as a frill on the dorsal surface (Fig. 1).

### Comparative diagnosis

Comparisons were made on the basis of reports of the following authors for concerned species: Beauchamp, 1930; Graff, 1899; Humbert and Glaparede, 1862; Moseley, 1878; Whitehouse, 1914, 1919; Wright, 1860 (Table 3). *Bipalium bengalensis* differs from its congeners by several characters, the most important of which is that it lacks any longitudinal band or line on the dorsal surface while in most of the species one (*B. dihangense*, *H. palnisia*), two (*H. dodabettae*), three (*B. splendens*, *B. andrewesi*) or five (*B. kewense*) bands or lines are present on the dorsal surface. Moreover, present species bears a mid-dorsal thin narrow groove (indistinct in live specimen) which is absent in all other species. Among 29 Indian species, only five species viz. *B. indicum*, *B. grayia*, *B. splendens*, *H. proserpina* and *B. smithi* are reported from West Bengal. Like *B. bengalensis*, *B. splendens* and *H. proserpina* are black in colour but *B. splendens* have three black lines while in *H. proserpina*, a wide black band is present on the dorsal surface. The present species differs from other three Bengal species by colour pattern (black vs. other than black).

In *B. bengalensis*, location of mouth and gonopore from anterior end is longer than those of *H. proserpina* (55.22 vs. 44.01% BL and 70.55 vs. 60.00% BL respectively). Compared to *B. smithi*, head breadth and position of gonopore length is shorter in the present species (15.56 vs.10.94% BL and 80.00 vs. 70.55% BL respectively). Distance of gonopore from anterior end is longer in *B. bengalensis* compared to that of *B. splendens* (70.55 vs. 60.00% BL) and sole is also wider to that of later species (27.01 vs. 22.22% BW). In *B. indicum*, though other two measurements like body width and position of gonopore from anterior end are within the range with those of *B. bengalensis* but position of mouth from anterior is shorter in the former species (50.00 vs. 55.22% BL). Moreover, the former species differs from the later by the presence of eyes in the neck.

### Etymology

Latin name, genitive case, meaning “bengal” in reference to its occurrence.

### Distribution and habitat

*Bipalium bengalensis* is known for the type locality at Suri in West Bengal. Suri is a small town of Birbhum district, West Bengal, India. The specimens were collected from the road side of Vivekanandapally (a locality of Suri). Thick vegetation is present on both sides of the road with a few shrub like *Calotropis procera*, *Cassia sophera* etc. On both sides open drain and stagnant water provide decomposed organic matter and water to the entire vegetation throughout the year. This supports rich population of soil animals including the present Bipaliid species. Moreover, organic matters like household leftovers and faecal substance of domestic animal like cow and goat enrich the soil regularly. At this place, other soil animals like different species of earthworm, millipedes, coleopteran larva, termites, isopods, ants and their larva, spiders are common. These provide food to the population of *Bipalium bengalensis*. There are other solid wastes like stones, bricks, debris etc. which provide shelter to different soil animals including the present species.

### Revision

Morphology and distribution of 29 Indian Bipaliid species are listed in Table 2. Morphological characters of these species in percentage of body length and body width are presented in Table 3. Besides these 29 species, Kawakatsu and Jayashankar (2013) reported three species of *Diversibipalium* (=*Bipalium*), one from Madurai and two from Bangalore. In all three species only length and width are mentioned. Although colour, shape and banding patterns were described vividly. *Bipalium kewense* was reported from Delhi state in the name *Placocephalus kewense* by Johri in 1952 and later Sharma and Sharma (1977) reported the same species from India. Patra and Aditya (2001) studied regeneration pattern of an unidentified land planarian from Darjeeling (West Bengal). The species is very long (15-20 cm) and wide (1-3 cm) with 3-5 prominent yellow or black stripe on the body.

**Table 2.**
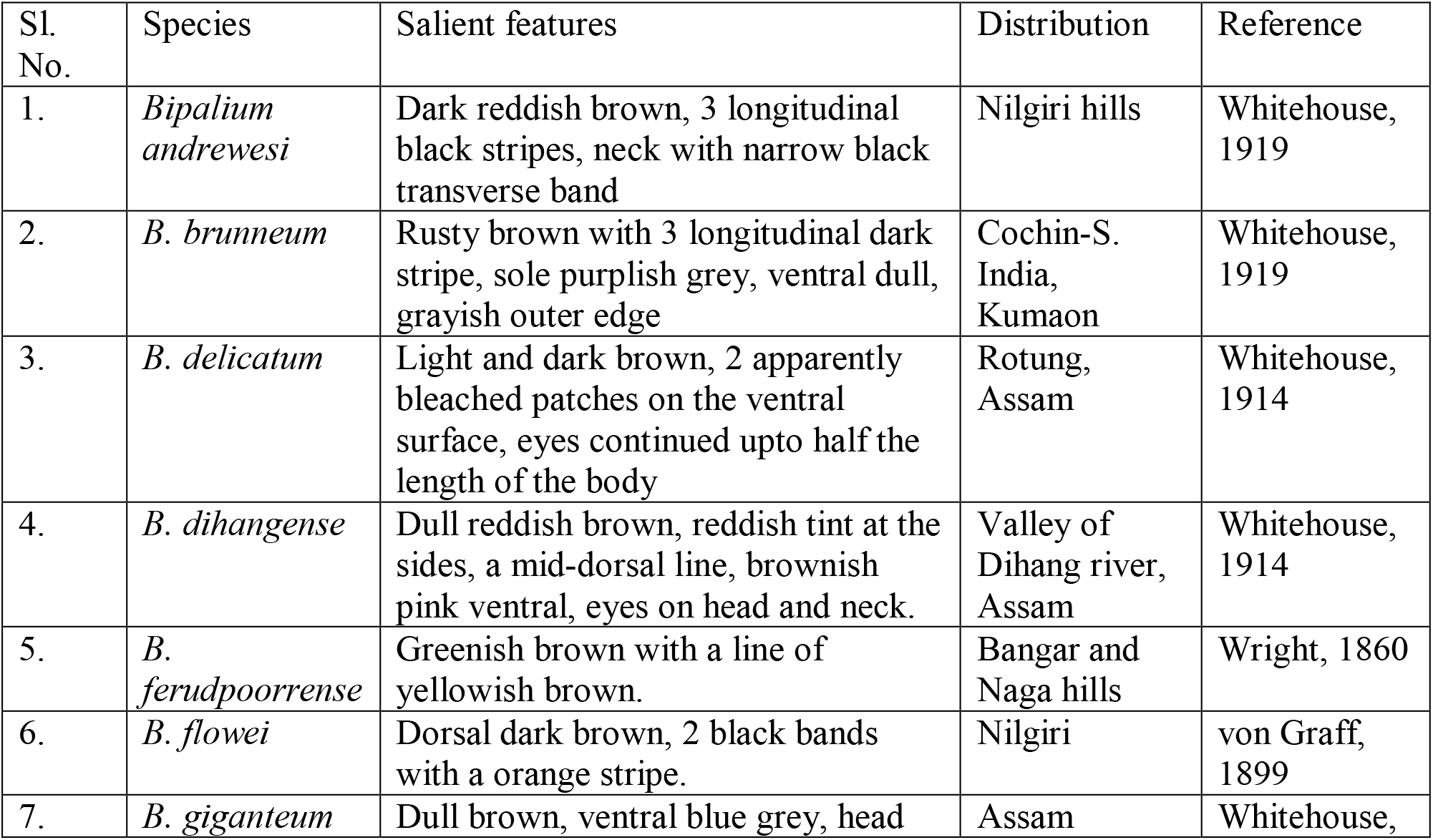
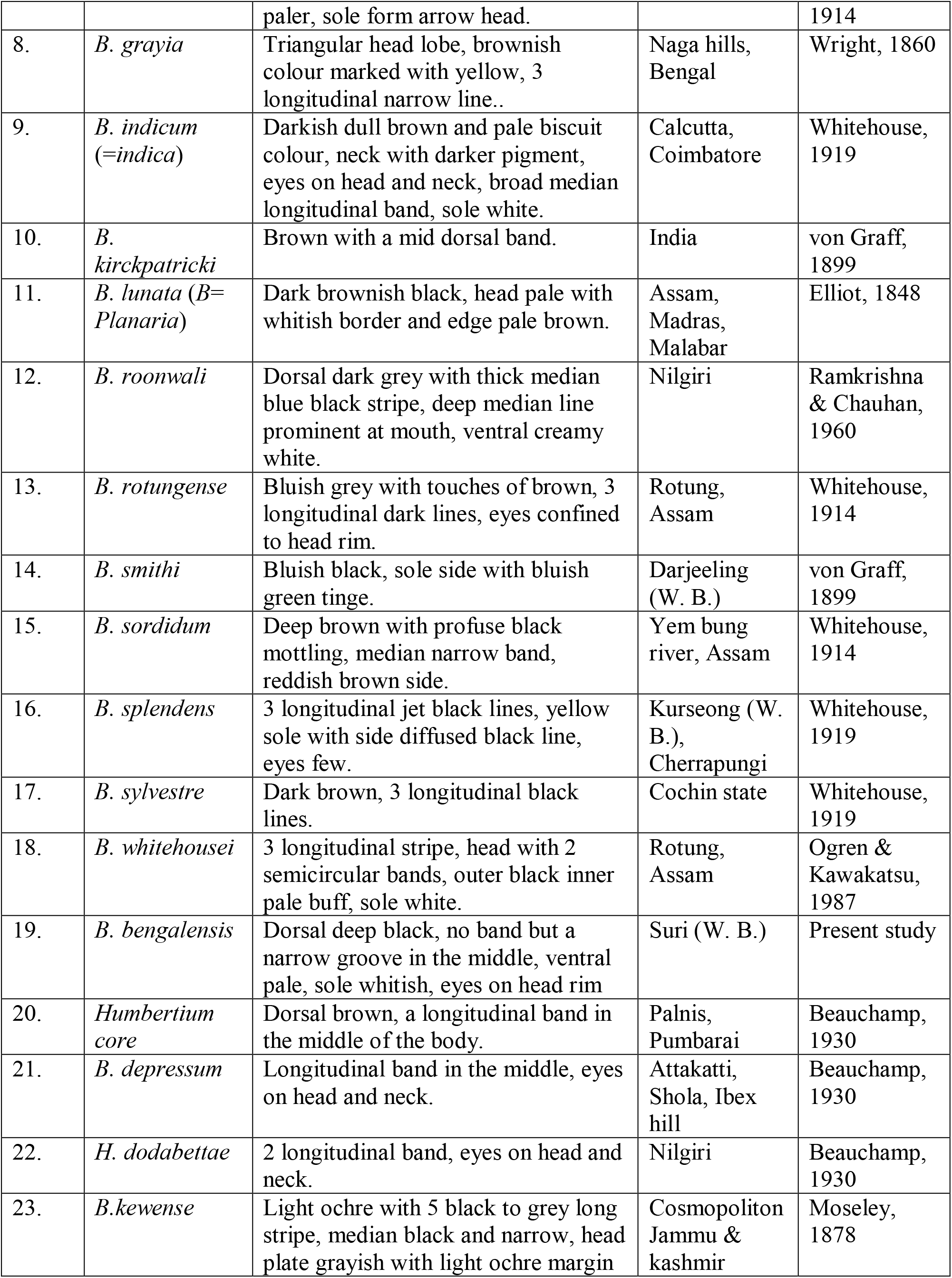
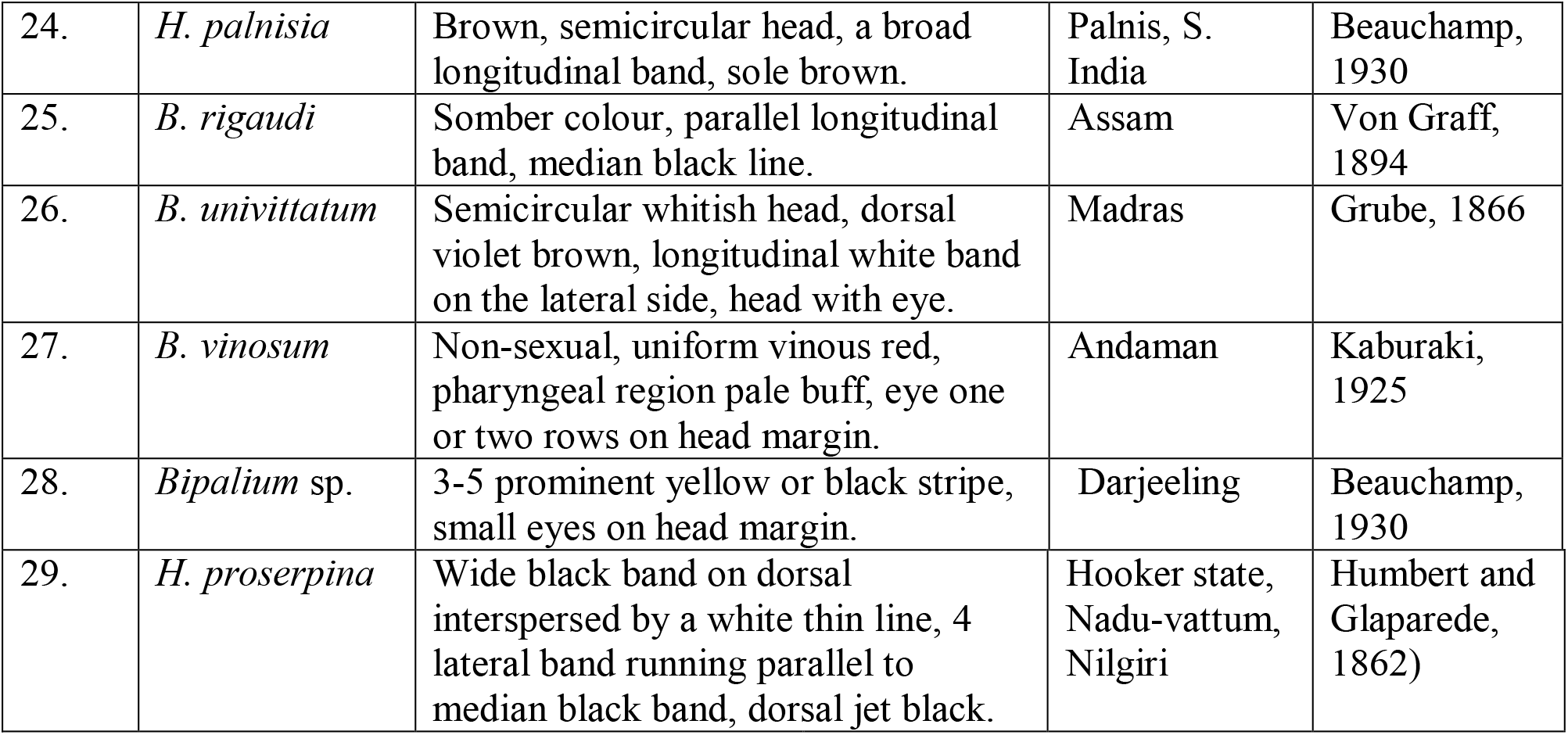
List of Indian Bipaliid species with their salient features and distribution in India.

**Table 3.**
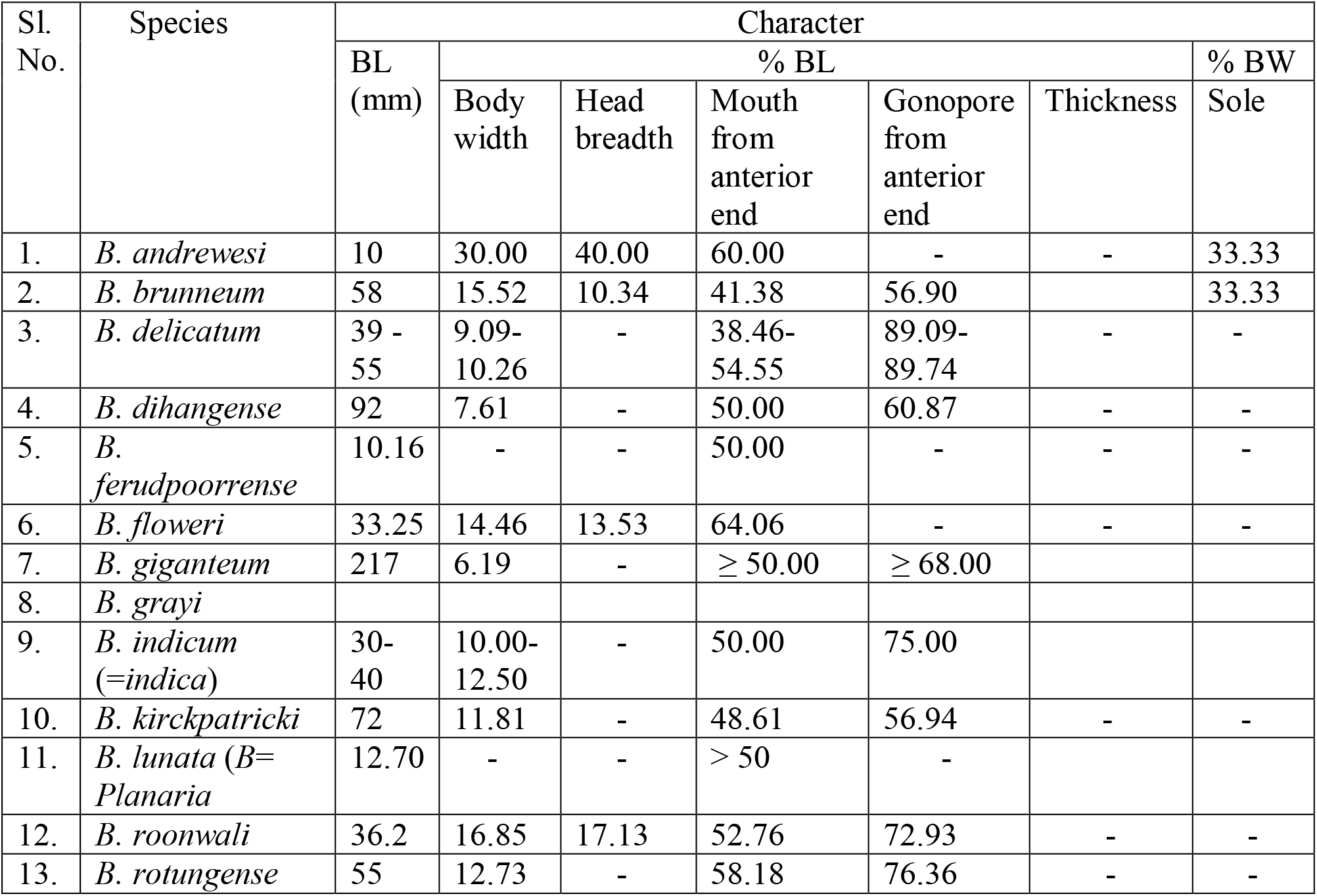
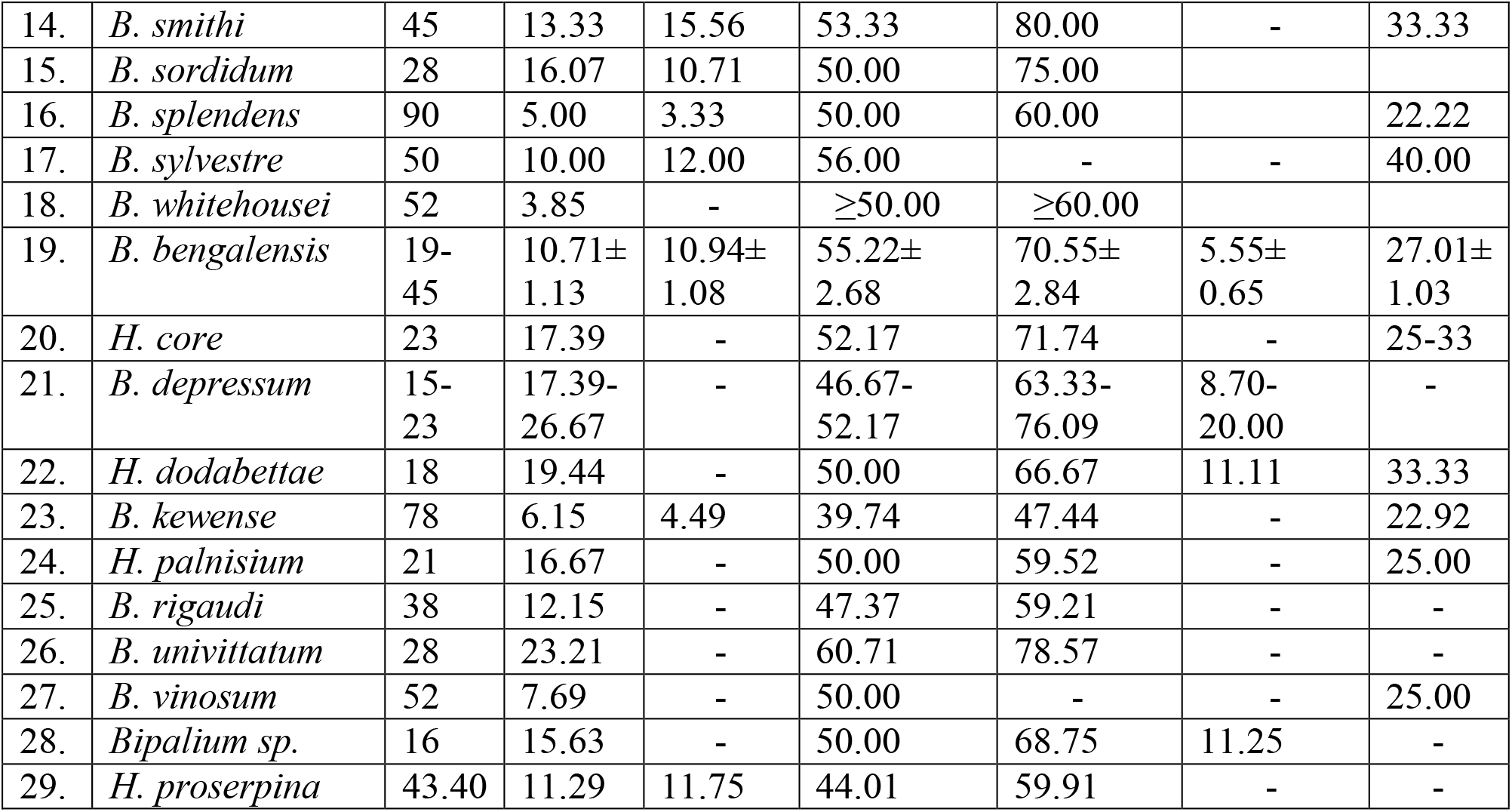
Biometric data of Indian Bipaliid species.

## Discussion

In practice, while describing a species of land planarian, author always put the actual or raw data of different body parts of an individual. Then it becomes very difficult to separate different species of unequal body length on the basis of size or position of body parts. To overcome this problem, it is better to present the data related to different body parts in proportion to body length or width as because in all the symmetrical animals, size of any body part is always proportional to its body length or width. Then it will be very easy to identify different species biometrically as shown in Table 3.

Except a few (the invasive *B. kewense*), most of the species are restricted to a particular geographic area as they are less dispersed. The land planarians are stenohygric in nature (Froehlich, 1955; Winsor et al., 1998; Sluys, 1999) and feed on a wide range of invertebrates like earthworm, isopods, insect larvae, termites etc. (Du Bois Reymond Marcus, 1951; Jones et al., 1995; Ogren, 1995; Sluys, 1999). Boag et al. (1990) commented that soil moisture content, temperature and availability of food are the principal factors to determine the presence of terrestrial planarians. All the above mentioned factors are available in the present habitat for survival of a large population of *B. bengalensis* for a long time. On the basis of morphological characters, biometric data and comparison with other species, it can be concluded that *B. bengalensis* is distinctly separated from its congeners of West Bengal and should be treated as a new species of the genus *Bipalium*.

So far the criteria of selection of a genus in Bipaliidae indicate that there are two genera, *Bipalium* and *Humbertium*, which can be distinguished clearly by one or more points. Another genus, *Diversibipalium* was created from *Bipalium* which lack information on sexual structure though each species is separated from each other morphologically. Winsor (1983) commented that Stimpson’s generic diagnosis which is based on external morphology proves to be unsatisfactory. But the argument is that gene regulates the morphology of an individual. So morphology based taxonomy should not be discarded rather it can further be verified by molecular taxonomy. The creation of a new genus (from *Bipalium* to *Diversibipalium*) on the basis of its anatomical description is not acceptable because morphologically described species after molecular taxonomy ascribed as synonymous or grouped into a species group as found in many other cases. Recently, molecular study of Mazza et al. (2016) indicates that *Diversibipalium multineatum* is a kin to *Bipalium nobile* and *Novibipalium venosum* appears to be a member of *Bipalium* species group. So it can be concluded that all the following genera are synonymous of *Bipalium*:

*Sphyrocephalus* Bleeker, 1844; *Planaria* Elliot,1848; *Dunlopea* Wright, 1860; *Sphaerocephalus* Loman, 1888; *Perocephalus* Graff, 1896; *Placocephalus* Graff, 1896; *Novibipalium* Kawakatsu et al., 1998 and *Diversibipalium* Kawakatsu, 2002.

## Acknowledgement

I am deeply indebted to my son Dr. Soumendranath Bhakat of Lund University, to take interest on the species and my wife for her continuous inspiration in this critical period of lockdown. I am also grateful to my colleagues of Rampurhat College for their cooperation and Mr. Sourav Saha of Satyajit Roy Film and Television Institute, Kolkata for technical help.

